# Hydrophobicity drives receptor-mediated uptake of heat-processed proteins by THP-1 macrophages and dendritic cells, but not cytokine responses

**DOI:** 10.1101/2020.07.02.184317

**Authors:** Ying Deng, Coen Govers, Malgorzata Teodorowicz, Ieva Liobyte, Ilaria De Simone, Kasper Hettinga, Harry J. Wichers

## Abstract

Impact of processing on immunogenicity of food proteins has clearly been demonstrated, but the underlying mechanisms are still unclear. In our previous study, the uptake of the cow’s milk protein β-lactoglobulin (BLG) by THP-1 macrophages varied after applying different processing methods and was positively correlated with hydrophobicity and aggregation. Here we applied the same 3 processing methods: wet heating (60 °C) and low- or high-temperature (50 °C or 130 °C, respectively) dry heating in absence or presence of reducing sugars (i.e. glucose, lactose or galacto-oligosaccharide) to lysozyme and thyroglobulin, which have different pI or molecular weight compared to BLG, respectively. Uptake of the soluble fraction was tested in two types of, genetically homogeneous, antigen-presenting cells (macrophages and dendritic cells derived from THP-1 monocytes). This revealed a strong correlation between the uptake of the different protein samples by macrophages and dendritic cells, and confirmed the key role of hydrophobicity, over aggregation, in determining the uptake. Several uptake routes were shown to contribute to the uptake of BLG by macrophages. However, cytokine responses following exposure of macrophages to BLG samples were not related to the levels of uptake. Heat-treatment-mediated increases in uptake did thus not induce a response in our read-out systems. Together, our results demonstrate that heat-treatment-induced increased hydrophobicity is the prime driving factor in uptake, but not in cytokine production, by THP-1 macrophages.

## 1. Introduction

Dietary proteins play an important physiological role, not only in providing amino acids and energy, but also for the maturation and regulation of the immune system. Proteins are known to be involved in the regulation of chronic inflammation and be the cause of allergies [1]. For example, a higher intake of animal protein was found to enhance the pro-inflammatory response of macrophages in mice [2]. Most of the proteins that humans ingest have been through a heating process as part of the food processing, as a result of which structural modifications, such as glycation (with reducing sugar) and/or aggregation of proteins, may occur.

In an earlier study on β-lactoglobulin (BLG), the major whey protein of milk [3], we observed that heat processing under different conditions altered the physicochemical properties and its uptake by THP-1 macrophages. In particular, heating in solution at 60 °C for 3 days introduced changes in hydrophobicity and molecular weight, which were shown to be the key determinants for uptake. This event may be relevant for a possible subsequent immune response [4, 5] and has been linked to heat-treatment-induced changes in protein’s propensity to induce an allergic reaction [6-8]. Despite the new findings in that study, obvious limitations relate to the use of a single protein, cell phenotype and immunological read-out.

Here, we extend previous findings by investigating 2 other proteins (lysozyme and thyroglobulin), another cell phenotype (THP-1 dendritic cells), and different immunological responses to yield novel insight. Lysozyme and thyroglobulin were subjected to the same heat treatments as used for BLG. Lysozyme from chicken egg white is a single chain polypeptide with four disulphide bridges and is a rather inflexible protein with stable structure and properties [9]. It has a molecular weight of 14.3 kDa which is similar to 18.4 kDa of BLG and an isoelectric point (pI) of 10.7 [10] which is clearly different from BLG (pI=5.2) [11], resulting in a positive net charge of this protein at the physiological pH. Moreover, it is also known as an allergen, as exemplified by the high incidence of sensitization and allergenicity to this protein [12]. On the other hand, bovine thyroglobulin, which originates from follicular cells of the thyroid gland, is a protein that is considered as less allergenic. It has a molecular weight of 330 kDa as a monomer, making it more than 18 times larger than the mass of the other two proteins under investigation. It has a complex quaternary structure composed of four subunits that are disulphide bonded [13, 14]. Its pI (4.6) is close to that of BLG, conferring it a similar charge as BLG at the pH of the exposure medium. We analysed uptake of heat-treated proteins by macrophages and dendritic cells and correlated this to physicochemical characteristics. Furthermore, we investigated the mechanism and receptors that are involved in uptake and identified downstream immunological responses by quantifying cytokine response.

Taken together, our findings provide insights into the role of protein size and pI towards physicochemical changes upon food-processing related heat-treatments. This appears to yield generalisable correlations between a protein’s physicochemical characteristics and its uptake by antigen-presenting cells (APCs). Finally, we identified routes of uptake and downstream immunological responses which will help to clarify the interaction of protein particles with macrophages and the potential immunoregulatory effect of it.

## 2. Materials and methods

### 2.1 Chemicals

All chemicals were purchased from Sigma Aldrich (St Louis, Missouri, USA) unless otherwise stated.

### 2.2 Sample preparation and fluorescent labelling

β-Lactoglobulin (BLG) from cow’s milk, lysozyme from chicken egg white and thyroglobulin from bovine thyroid were dissolved in sodium phosphate buffer (10 mM, pH 7.4, same below) to a protein concentration of 5 mg/mL. Besides this solution without saccharides, D-glucose (glu), D-lactose (lac), and galacto-oligosaccharide (GOS; Royal FrieslandCampina, Wageningen, the Netherlands) were added to reach a final 1:4 molar ratio of total free amino groups to saccharide reducing ends. Identical heating methods, being high-temperature dry-heating (H), wet-heating (W) and low-temperature dry-heating (L), sample preparation and naming were applied to the proteins as indicated in a previous study [3]. Fluorescein isothiocyanate isomer I (FITC) labelling was performed as described previously [3].

### 2.3 THP-1 cell culture and differentiation

The human monocytic leukaemia cell line THP-1 (ATCC, Manassas, Virginia, USA) was differentiated into cells characteristic for resting macrophages (M0) as described elsewhere [15], at a concentration of 1×10^6^ cells/mL using 100 ng/mL phorbol-12-myristate-13-acetate (PMA) for a 48 hours stimulation followed by washing two times with medium and another 48 hours of rest. Using a cell concentration of 0.25×10^6^ cells/mL and incubation with 20 ng/mL of IL-4 and 20 ng/mL of PMA for 4 days as described [16], immature dendritic cells were generated from THP-1 monocytes.

### 2.4 Physicochemical analysis

The analysis of solubility (protein mass remaining in solution), loss of amino group (OPA method), AGE formation (fluorescent measurement), exposure of hydrophobicity region (ANS method), surface charge (zeta-potential measurement), secondary structure (circular dichroism measurement) and aggregation (size exclusion chromatography) were performed using the protocols as described previously [3].

### 2.5 Uptake and blocking of different uptake routes

Uptake experiments were performed as described previously [3]. No significant difference of absolute uptake value has been found for native BLG, thyroglobulin or lysozyme in macrophages or dendritic cells (S1 Fig). As there is no bias for the basic uptake capability for the different proteins, all the uptake results were presented relative to their corresponding native protein to be able to compare among different proteins. To determine which route of uptake was involved in a protein’s uptake by THP-1-derived cells, the cells were pre-incubated for 30 minutes with DMSO (control) or different inhibitors: 25 µg/mL nystatin (caveolae-dependent uptake inhibitor), 10 µg/mL chlorpromazine (clathrin-dependent uptake inhibitor), 10 µg/mL cytochalasin B (microphagocytosis inhibitor) or all inhibitors combined, dissolved in DMSO. Following incubation, the cells were washed with PBS and incubated for 2 hours with FITC-labelled protein samples. Subsequently, the cells were harvested and measured using flow cytometry as described previously [3]. For all uptake experiments 5 µl trypan blue was added per 100 µl of cell suspension to quench extracellular FITC signal before flow cytometry analysis. Uptake was calculated as either the fold change in mean fluorescence intensity (MFI) relative to control, after correcting for FITC labelling efficiency (control = 1), or as percentage to control (control = 100%).

### 2.6 Soluble receptor for advanced glycation end products (sRAGE), CD36 and Galectin-3 inhibition ELISA

The experiment was performed according to the method of Liu et al. [17]. As a positive control, soy protein (Bulk Powder, Colchester, UK) was mixed with D-glucose in an equal w/w ratio in 10 mM PBS buffer (pH 7.4), to a protein concentration of 10 mg/mL and heated for 90 minutes at 120 °C. The positive control was diluted in 1.5 mM of sodium carbonate buffer pH 9.6 to 20 µg/ml and used for plate coating by adding 100 µL per well to a Nunc MaxiSorp(tm) flat-bottom plate (Thermo Fisher, Waltham, Massachusetts, USA) and incubating overnight at 4 °C. Then the plate was washed 3 times with PBS buffer pH 7.4 containing 0.05% (v/v) Tween-20. After blocking with 3% bovine serum albumin (BSA) in PBS for 1 hour at room temperature, the wells were washed 3 times and incubated with 80 µL protein samples (25 μg/mL) at 37 °C for 1 hour. The samples were preheated at 37 °C for 45 minutes with recombinant human sRAGE, CD36 or galectin-3 at a concentration of 1, 0.5 or 3 µg/mL (R&D Systems, Minneapolis, Minnesota, United States) in PBS buffer with 1.5% BSA and 0.025% Tween-20. For detection, the plate was washed 3 times and incubated with 80 µL per well of monoclonal mouse IgG2B human sRAGE or galectin-3 antibody (R&D Systems, Minneapolis, Minnesota, United States) at a concentration of 0.2 µg/ml for 30 minutes shaking at room temperature. After washing 4 times the plate was incubated for 30 minutes while shaking at room temperature with 80 µL per well polyclonal goat anti-mouse HRP-conjugated (DAKO, Glostrup, Denmark) at a concentration 0.25 µg/mL. For galectin-3, an additional incubation step with 0.2 µg/ml Streptavidin (SDT, Baesweiler, Germany) 80 µL per well for 20 minutes was followed. For CD36, only incubation with goat anti-human IgG/HRP (SouthernBiotech, Birmingham, Alabama, United States) at a concentration of 0.25 µg/ml for 30 minutes shaking at room temperature was used for the detection step. For all receptor inhibition ELISAs, the plate was then washed 4 times and 80 µL per well TMB substrate was added and incubated for 10 minutes before stopping the reaction by adding 100 uL of 2% HCl per well. The absorbance measured at 620 nm by an Infinite® 200 PRO NanoQuant (Tecan, Männedorf, Switzerland) was subtracted from the measured absorbance at 450 nm. Each sample was measured in triplicate and values were averaged. Absorbance of buffer was used as a control and was subtracted from every sample. The absorbance of the control solution without any inhibition was coded by Abs_control._ The percentage of inhibition for samples was equal to (Abs_control_-Abs_sample_)/Abs_control_*100%.

### 2.7 Cytokine production measurement

THP-1 macrophages were incubated with 100 μg/mL of protein sample or medium as control for 16 hours, after which the supernatant was collected. The cytokine production was measured using the ELISA Deluxe Set Human IL-8/CCL20/IL-1β (Biolegend, San Diego, California, United States), following the manufacturer’s protocol.

### 2.8 Statistics

The statistical analysis was performed using Prism 6 software (GraphPad Software, San Diego, California, United States) with p < 0.05 considered to be significant. The correlation analysis was done by calculating the Pearson correlation coefficient (r) and the two-tailed p value. The PCA plot was generated with the dataset of variables after being mean centred and weighted by 1/standard deviation using Unscrambler software (CAMO, Oslo, Norway).

## 3. Results

### 3.1 Wet-heating significantly increased uptake of thyroglobulin by THP-1-derived macrophages whereas heat-treatment did not increase uptake of lysozyme

Thyroglobulin and lysozyme were wet-heated (W), high-temperature dry-heated (H) or low-temperature dry-heated (L) in the absence or presence of saccharides with different lengths: monosaccharide (glucose), disaccharide (lactose) or oligosaccharide (GOS). Soluble protein from each sample was concentration calibrated, fluorescently labelled and incubated with resting macrophages that were derived from THP-1 monocytes. High-temperature dry-heating of thyroglobulin in the presence of glucose and lactose and lysozyme in the presence of glucose led to complete insolubility (S2A and S4A Fig) and therefore these samples were excluded from further analysis. Prior to the experiment, the lipopolysaccharide (LPS) contamination was measured (Table in S7 Table) and found to be below the threshold level that significantly affected uptake capacity of THP-1 macrophages (i.e. 1000 ng/mL), as reported in an earlier study [3].

For uptake of thyroglobulin samples by macrophages (Fig 1A), the type of heat treatment appeared to play a more determining role than the presence or absence of saccharides. Notably, all wet-heated samples, regardless of the presence of saccharides, showed significantly increased uptake compared to untreated protein. Although not significantly differing from untreated protein, high- and low-temperature dry-heating increased the uptake of thyroglobulin as well. On the contrary, no significant differences in uptake of lysozyme after different treatments and saccharide exposures was noticeable (Fig 1B).

**Fig 1.**
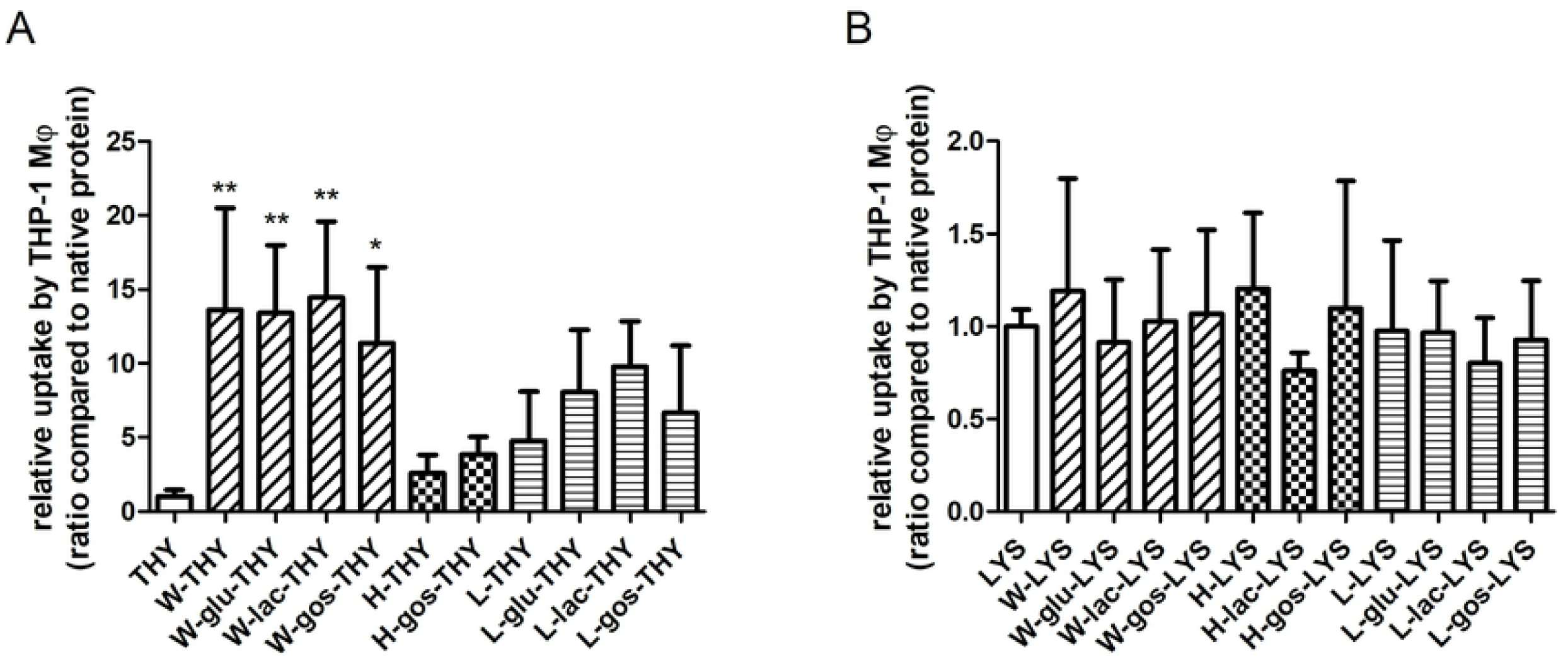
Uptake of thyroglobulin by macrophages was significantly increased upon wet-heating of the sample. Thyroglobulin (THY; A) and lysozyme (LYS; B) were untreated or heated (W, H or L) in the absence or presence of saccharides (glu, lac or GOS). THP-1-derived macrophages (Mϕ) were incubated with FITC-labelled soluble fraction of protein samples for 2 hours after which uptake was determined using flow cytometry. The results represent mean values ± SD of n=4 measurement of 2 independent experiments based on 2 independent sample sets. Statistical differences compared to native proteins were calculated with Dunnett’s Test: *p < 0.05; **p < 0.01.

### 3.2 Uptake of heat-treated protein samples by THP-1-derived immature dendritic cells is qualitatively similar to that of THP-1-derived macrophages

The uptake of processed thyroglobulin and lysozyme, as well as BLG which was previously reported [3] was only tested for macrophages. To expand the understanding of the proteins’ uptake beyond only macrophages, THP-1-derived immature dendritic cells (iDC) were also tested, with identical methodology as for macrophages. The uptake of wet-heated thyroglobulin in the absence or presence of saccharides by these iDC was significantly increased compared to the native protein (Fig 2A). For lysozyme, the relative uptake remained the same regardless of the treatment (Fig 2B). The uptake of wet-heated BLG in the absence or presence of saccharides by iDCs was also increased compared to the native form, although only reaching significance when heat-treated in the presence of glucose (Fig 2C). When compared to the results obtained with macrophages, the increase in uptake upon protein wet-heating for iDC was less pronounced (BLG-macrophage data from previous study [3]). However, the uptake pattern of all samples correlated strongly (r > 0.8, p < 0.0001) between the two cell types (Fig 2D).

**Fig 2.**
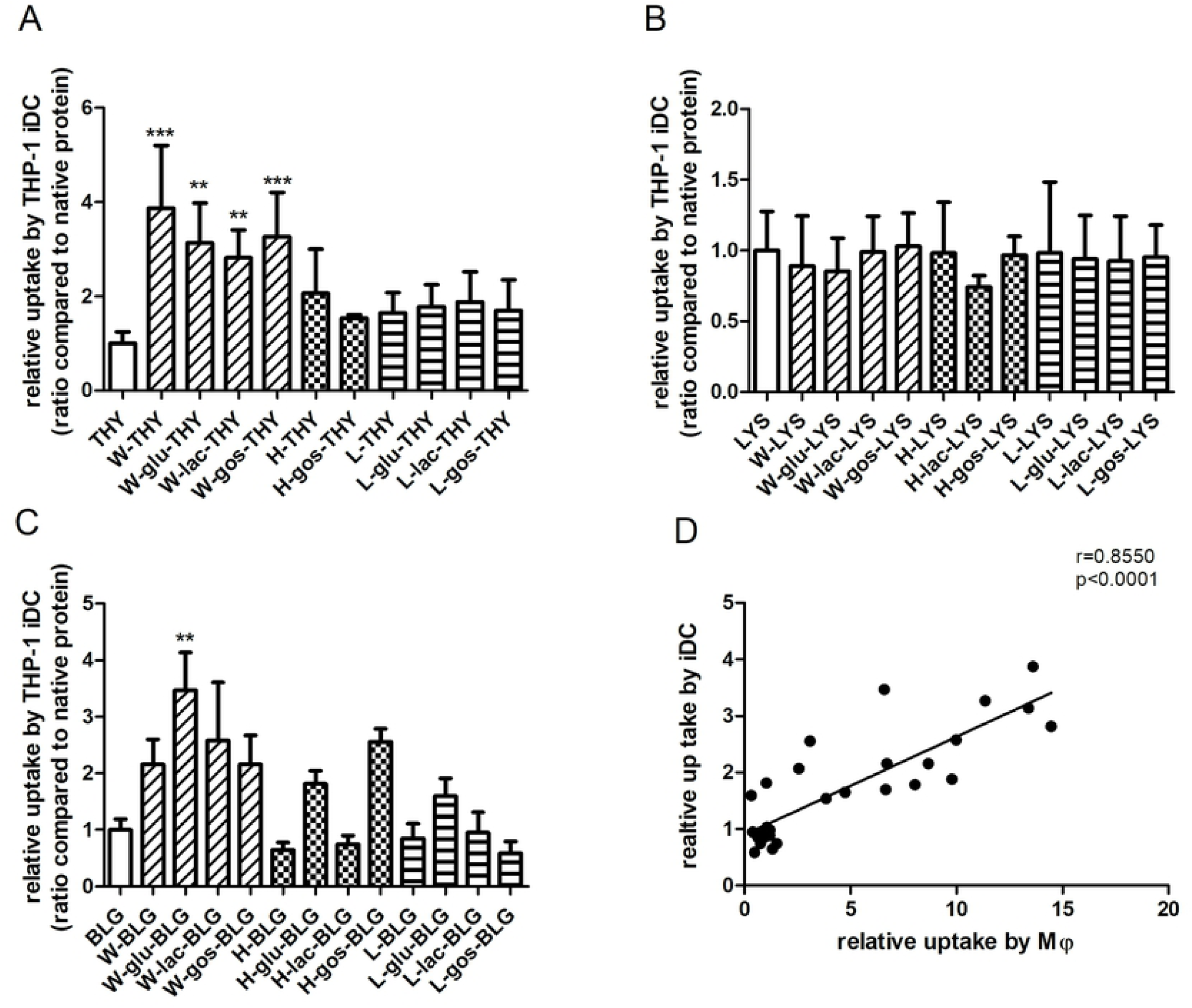
Uptake of thyroglobulin, lysozyme and BLG samples by THP-1-derived iDC was strongly and significantly correlated with uptake by macrophages. (A-C) Thyroglobulin (THY), lysozyme (LYS) and BLG were untreated or heated (W, H or L) in the absence or presence of saccharides (glu, lac or GOS). THP-1-derived iDC were incubated with FITC-labelled soluble fraction of protein samples for 2 hours after which uptake was determined using flow cytometry. The results represent the mean values ± SD of n=4 measurement of 2 independent experiments based on 2 independent sample sets. Statistical differences compared to native proteins were calculated with Dunnett’s Test: **p < 0.01; ***p < 0.001. (D) The relative uptake values of all protein samples by iDC was plotted against macrophages (Mϕ) and the correlation analysis was done by calculating the Pearson correlation coefficient (r) and two-tailed p value.

### 3.3 Increased uptake of thyroglobulin by THP-1-derived macrophages was correlated to hydrophobicity and aggregation

Due to the strongly correlated responses of iDC and macrophages in uptake, with a higher relative uptake by macrophages, the latter was used for further testing. To explain the physicochemical mechanism behind the differing uptake capability of macrophages for differently treated thyroglobulin and lysozyme, a set of measurements was performed on the soluble fraction after treatment.

For thyroglobulin, dry-heating at high-temperature significantly decreased its solubility, resulting in the absence of a soluble fraction when heating was performed in the presence of glucose and lactose (S2A Fig). The soluble fraction of samples in the presence of GOS or without saccharide revealed a strong decrease of free amino groups and increases in AGE-related fluorescence and formation of polymers in the presence of GOS (S2B, S2C and S3 Fig). Wet-heating did not result in loss of solubility, but also led to formation of polymers and, albeit less pronounced as for high-temperature dry-heating significant, losses of amino groups and increases in AGE-related fluorescence. Moreover, wet-heating induced a significant increase in the exposure of hydrophobic regions as measured by ANS-binding (S2D Fig). Dry-heating at low-temperature did not have much influence on the measured parameters for thyroglobulin, except for a significant loss of amino groups and a slight increase in oligomer and polymer formation (S2B and S3 Fig). There was no significant change in the secondary structure for any of the treated thyroglobulin samples (S2E and S2F Fig).

Applying identical heating and glycation conditions as for thyroglobulin did not generate any soluble aggregates for lysozyme (size exclusion chromatography, data not shown). High-temperature dry-heating led to a significant decrease of solubility with complete insolubility for the samples heated in the presence of glucose (S4A Fig). Among the soluble fraction of the high-temperature dry-heated samples, there was a significant decrease of free amino groups, increases of AGE-related fluorescence and exposure of hydrophobic regions in the presence of saccharides (S4B-D Fig). Moreover, results showed a significant loss of β-sheet structure for all samples except when heated in the presence of GOS (S4F Fig). Similarly, low-temperature dry-heating resulted in a significant loss of β-sheets in all lysozyme samples. For wet-heated lysozyme, only the exposure of hydrophobic regions in the presence of saccharides was significantly increased (S4D Fig).

To identify the most important physicochemical properties of heat-processed thyroglobulin that are related to its uptake by macrophages, we performed a principle component analysis (PCA) and a correlation analysis. In the PCA plot (Fig 3). All low-temperature dry-heated samples clustered together with the untreated thyroglobulin, indicating that the physicochemical properties of these samples were similar. Uptake was strongly related with wet-heating and positively related to hydrophobicity (ANS) and polymer proportion, and inversely to monomer and oligomer proportion. These relations were further confirmed by correlation analyses. As shown in Fig 4, there was a strong correlation between uptake and hydrophobicity and a moderate correlation between uptake and aggregation, as indicated by the fraction of monomers, polymers and oligomers. Of note is that the samples were strongly clustering into two groups at the extremes of the data scale for both the uptake and monomer or polymer content correlation plots (Fig 4B and 4C); the correlation between uptake and oligomer content was less profound (Fig 4D, p > 0.05). Contrastingly to thyroglobulin, no clear correlation could be found between treatment and physicochemical properties for lysozyme in the PCA plot (S5 Fig).

**Fig 3.**
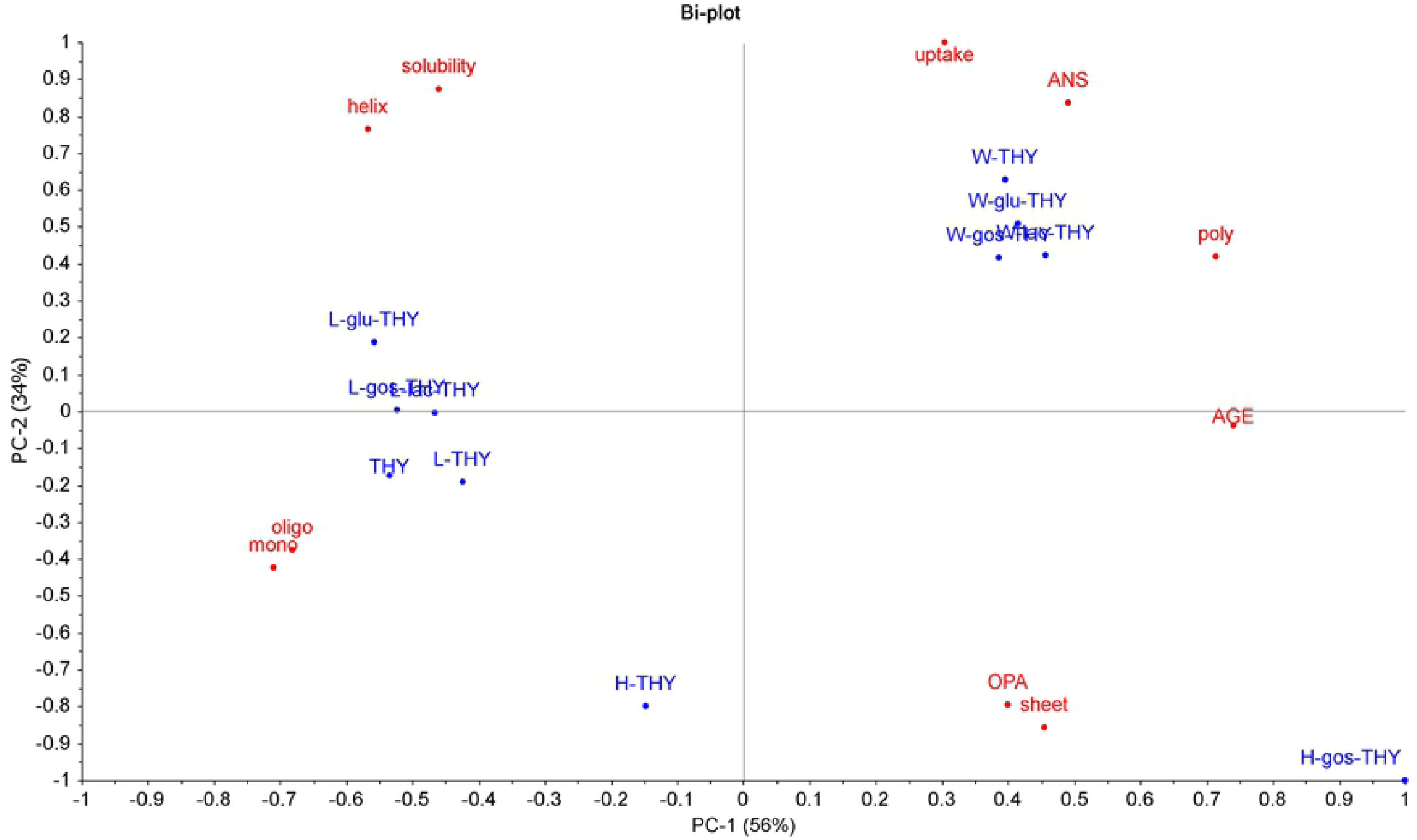
Uptake of thyroglobulin by macrophages is related to wet-heating, hydrophobicity and aggregation. Thyroglobulin (THY) was untreated or heated (W, H or L) in the absence or presence of saccharides (glu, lac or GOS) and tested for solubility, uptake of soluble fraction by THP-1 macrophages (uptake), AGE formation (AGE), glycation (OPA), percentage of α-helix (helix) or β-sheet (sheet), aggregation (monomer and smaller (mono), oligomers (oligo), polymers (poly)) and exposure of hydrophobic regions (ANS). PCA scores of the samples were given in blue and the parameter loadings of the principal components were given in red.

**Fig 4.**
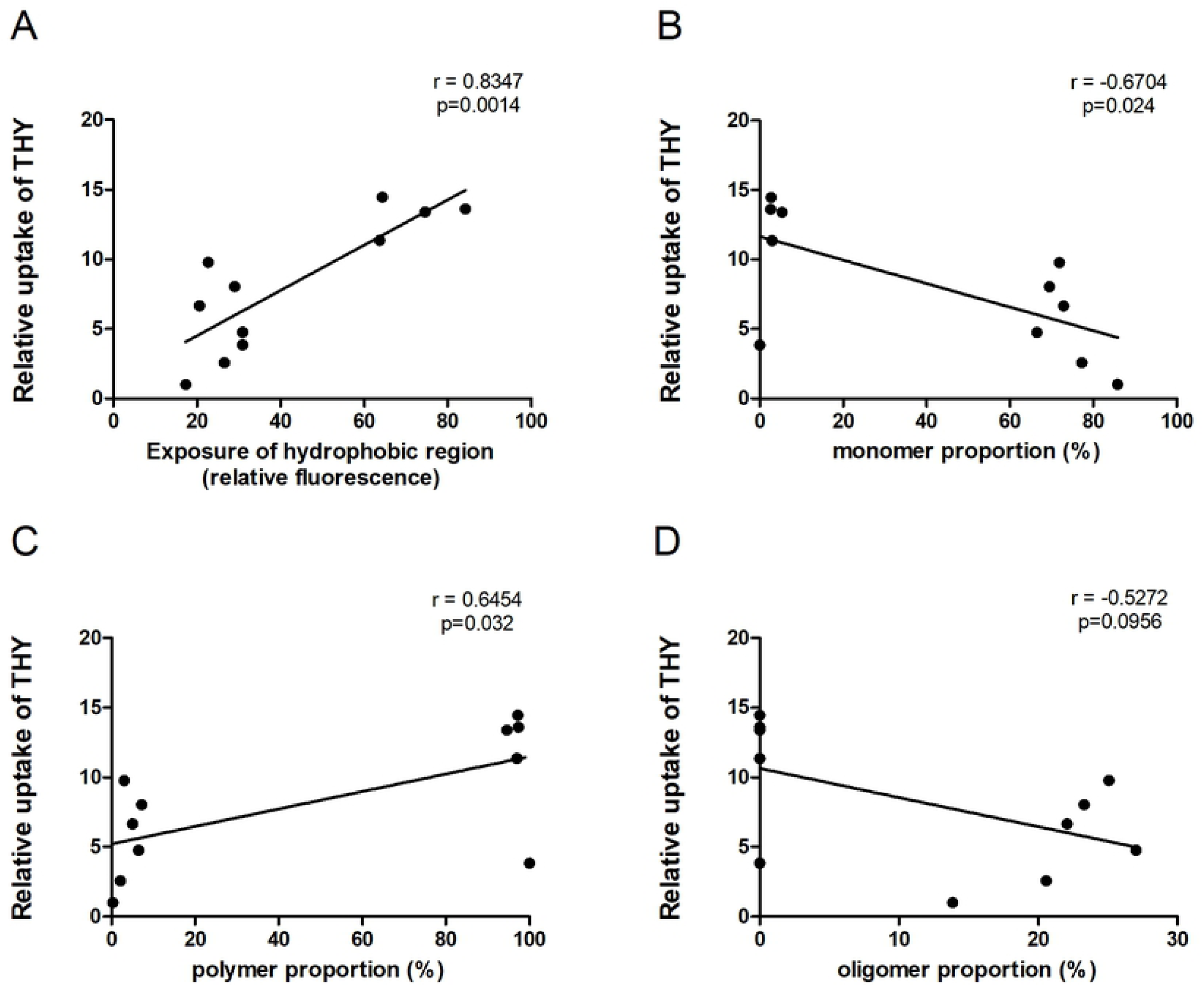
Correlation analysis of uptake of heat-treated thyroglobulin with physicochemical parameters. Uptake of soluble fraction of thyroglobulin (THY) samples by macrophages was correlated to hydrophobicity (|r|>0.7) and proportion of monomer, polymer and oligomer (0.7>|r|>0.5). The correlation analysis was done by calculating the Pearson correlation coefficient (r) and two-tailed p value.

### 3.4 EDC-mediated increase in hydrophobicity, but not cross-linking, is the main factor driving uptake by macrophages

EDC links the carboxyl and amino group of amino acids, and thereby offers the possibility to crosslink proteins without heating. This allows for a discrimination between aggregation and other heating-related effects, like hydrophobicity changes, with regard to their impact on uptake. As shown in Fig 5A and 5B, the proportion of polymer and monomer significantly increased, resp. decreased, in the EDC cross-linked BLG and thyroglobulin. On the contrary, only limited aggregation of lysozyme was observed upon EDC-treatment. EDC crosslinked BLG and thyroglobulin did not demonstrate an increase in uptake. Surprisingly, EDC-mediated crosslinking significantly increased the hydrophobicity of lysozyme (Fig 5C) and also its uptake by macrophages (Fig 5D).

**Fig 5.**
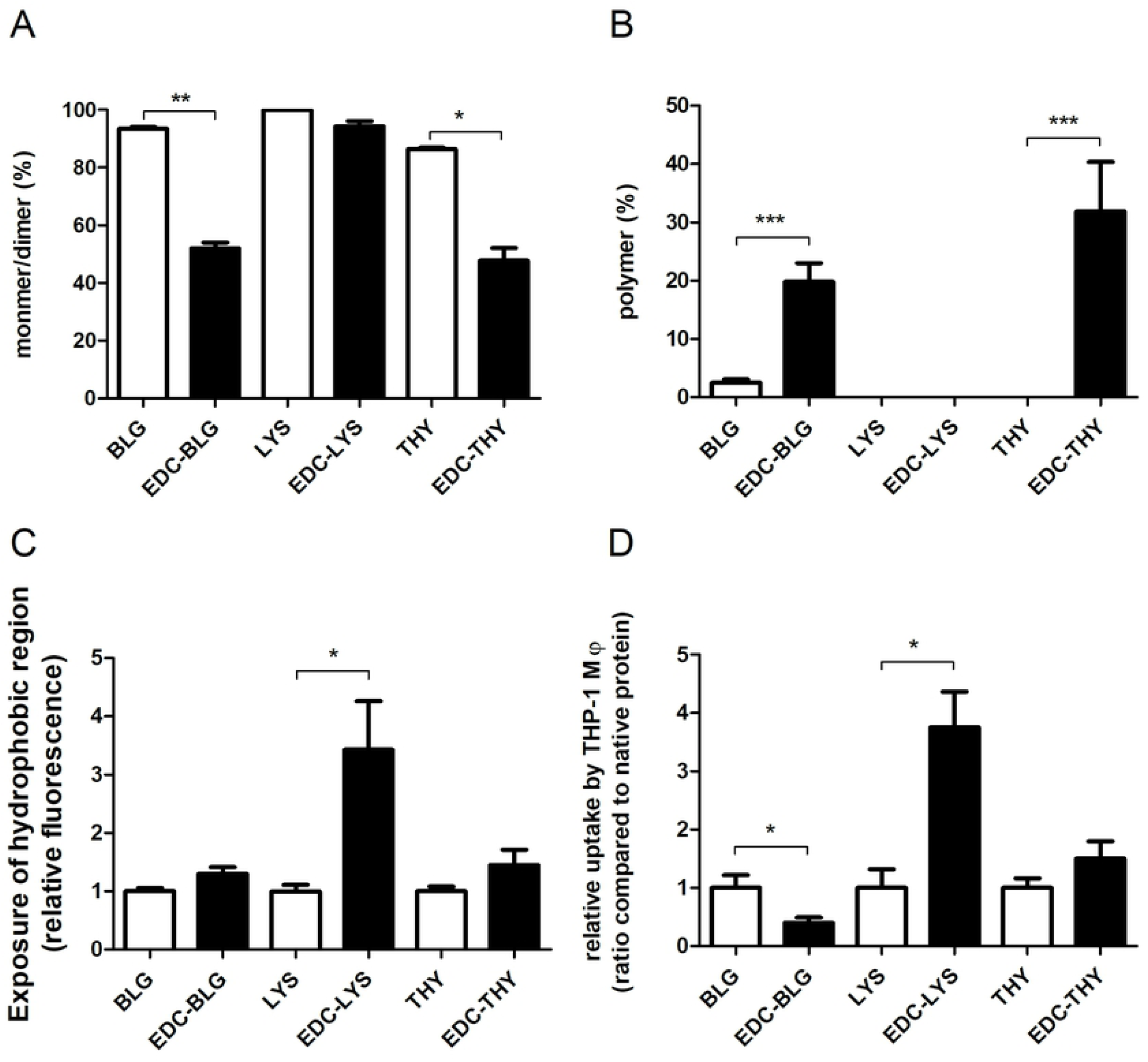
EDC-mediated hydrophobicity, but not aggregation, resulted in increased uptake of proteins by THP-1 macrophages. Thyroglobulin (THY), lysozyme (LYS) and BLG were non-treated or treated with EDC and subsequently analysed for the proportion of polymer (A), the proportion of monomer/dimer (B), and protein hydrophobicity (C). EDC-treated and non-treated proteins were fluorescently labelled and their uptake by macrophage (Mϕ) was measured (D). The results represent the mean values ± SD of 3-4 independent experiments with 2 independent sample sets. Statistical differences were calculated with two-tailed unpaired T-test with Welch’s correction: *p < 0.05; **p < 0.01; ***p < 0.001.

### 3.5 Uptake of heated-treated BLG was mediated via several routes

To better understand the mechanism of uptake, we set out to identify which routes of uptake were utilized for the various proteins that differed in physicochemical characteristics and therefore in their recognition motifs for cells. Native and heat-treated BLG in the presence of glucose was used as a subset, because these samples covered the whole range of levels of physicochemical modifications, and macrophages were pre-treated with inhibitors targeting phagocytosis, caveolae- or clathrin-mediated uptake or a combination thereof. Pre-incubation of macrophages with nystatin (caveolae inhibitor) and the combination of inhibitors, prior to BLG samples exposure, significantly inhibited the uptake (Fig 6). The usage of cytochalasin (microphagocytosis inhibitor) significantly decreased the uptake of both high-temperature dry-heated and wet-heated BLG (Fig 6B and 6C). Moreover, chlorpromazine significantly reduced uptake of wet-heated BLG, indicating that clathrin was involved in this process (Fig 6C).

**Fig 6.**
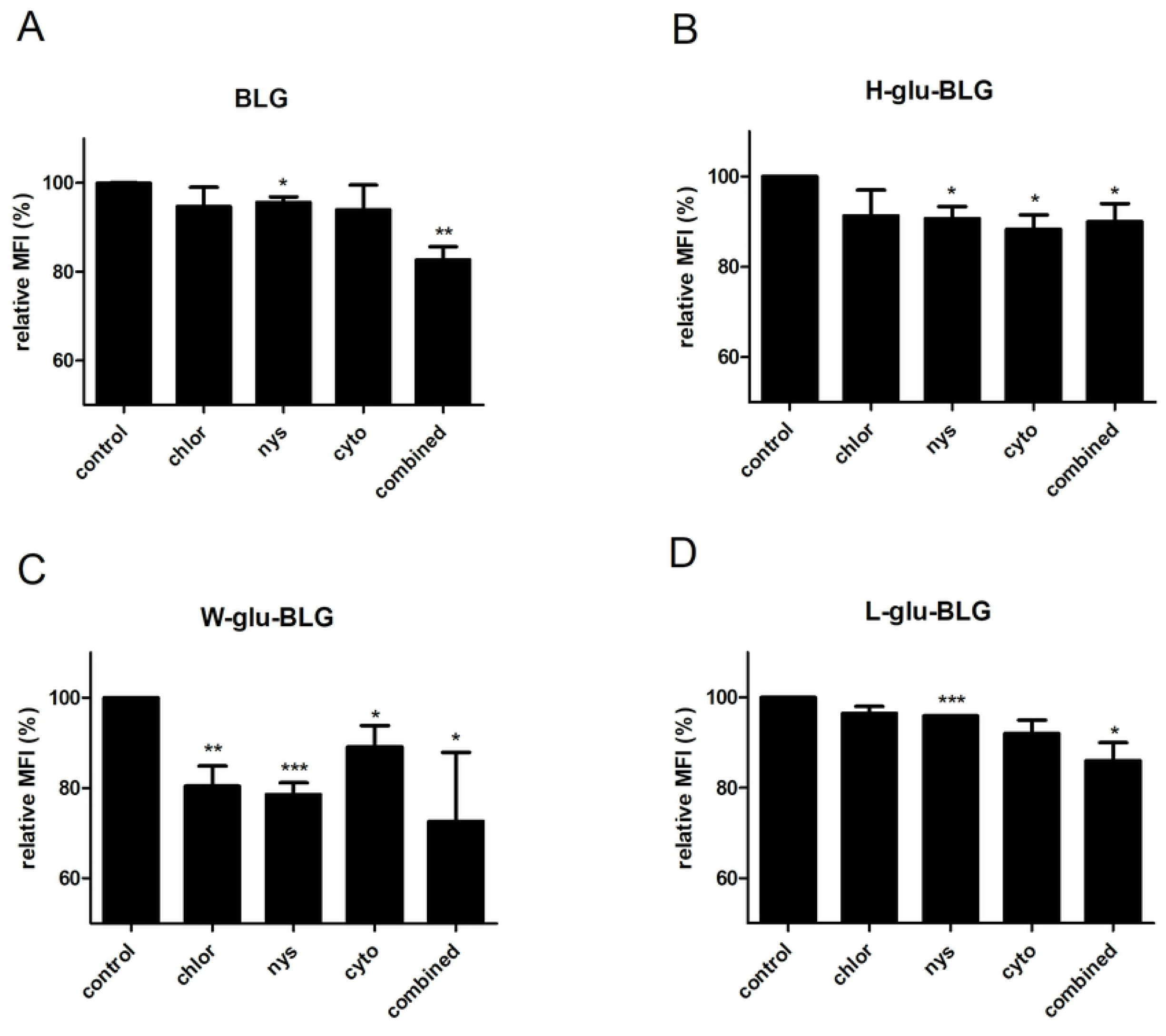
Several routes appeared to be involved in the uptake of wet-heated BLG. BLG was untreated or heated (W, H or L) in the presence of glucose (glu) and the soluble fraction was FITC-labelled. Macrophages were pre-treated with DMSO (control) or inhibitors (dissolved in DMSO) of clathrin-dependent uptake (chlorpromazine; chlor), caveolae-dependent uptake (nystatin; nys), microphagocytosis (cytochalasin; cyto) or a combination of them (combined) for 30 min before a 2-hour incubation with protein samples. Protein uptake was subsequently measured using flow cytometry. The relative MFI was calculated by dividing the MFI of a sample by the MFI of the corresponding control. The results represent mean values ± SD of multiple independent cell experiments (N=2-4) with 2 independent sample sets. The statistical differences were calculated with one-tailed unpaired T-test compared to control, *p < 0.05, **p < 0.01, ***p < 0.001.

### 3.6 Inhibition ELISA showed high binding of high-temperature dry-heated and wet-heated BLG by sRAGE, CD36 and galectin-3

We investigated several receptors that might be involved in binding and subsequent uptake of the BLG samples that were heat-treated in the presence of glucose. Using an inhibition ELISA, the binding of the BLG samples to the soluble receptor for advanced glycation end products (sRAGE), CD36-scavenger receptor and galectin-3 was analysed (Fig 7). Native BLG did not show any binding to sRAGE, galectin-3 or CD36. In contrast, high-temperature dry-heating of BLG induced significant receptor binding for all three tested receptors. Wet-heating of BLG similarly resulted in binding to all three receptors, albeit to a lesser extent and only significantly for sRAGE and galectin-3. Finally, low-temperature dry-heating did not induce any receptor-binding.

**Fig 7.**
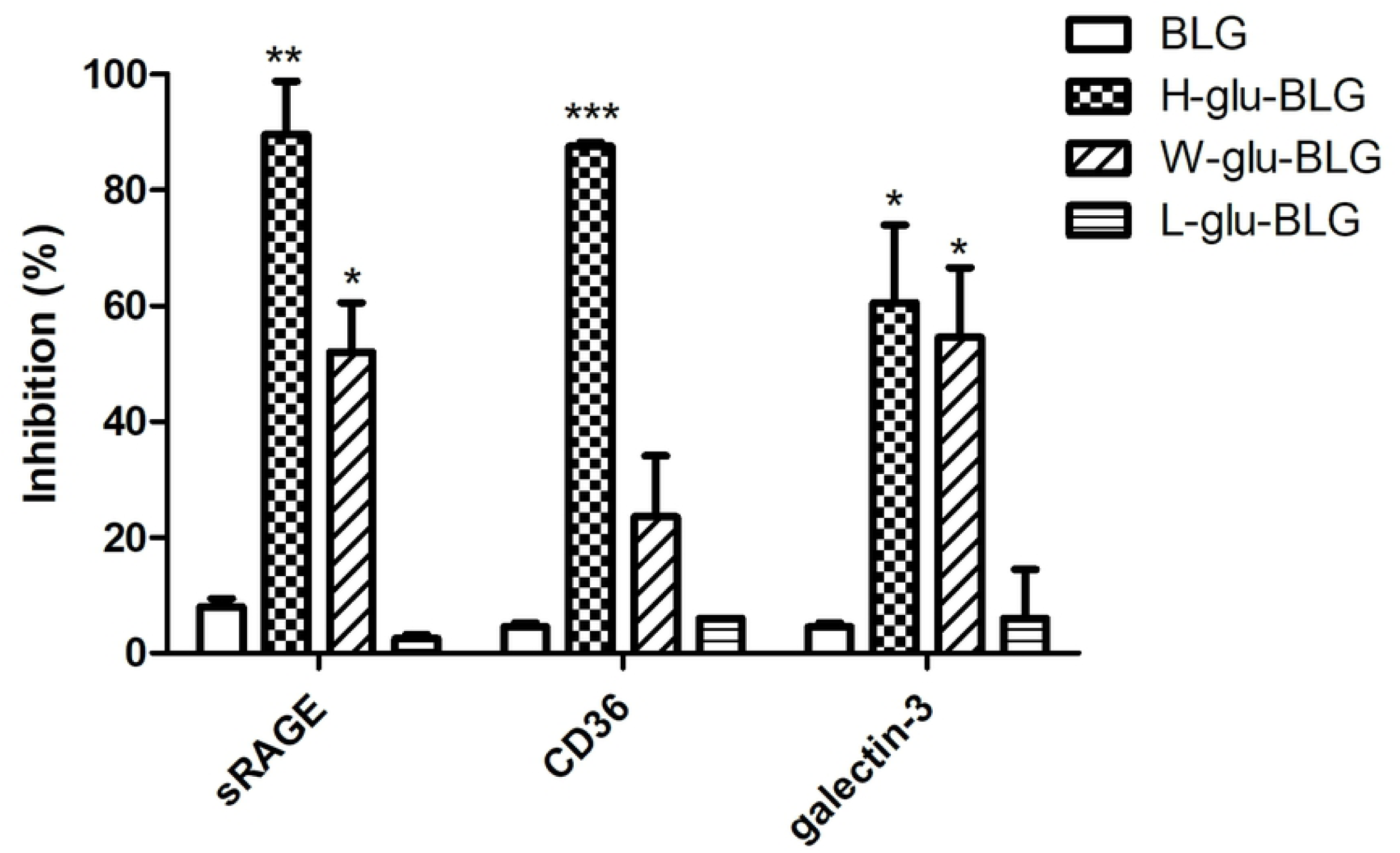
Heat-treatment in the presence of glucose confers receptor binding capacity to BLG. BLG was untreated or heated (W, H or L) in the presence of glucose (glu) and the soluble fraction was used to inhibit the binding of receptor to a positive control in the inhibition ELISA. Data shown are the mean values ± SD of n=4 based on measurement of 2 parallel prepared samples. Statistically significant differences between native and processed BLG for each receptor was determined using unpaired two-tailed t-Test, *p < 0.05, **p < 0.01, ***p < 0.001.

### 3.7 Cytokine responses of macrophages following exposure to BLG were reduced upon heat-treatment of BLG

Receptor binding and uptake of protein samples can lead to immune activity. The immunological response could be estimated by the expression of secreted cytokines or chemokines. Upon exposure to medium or native or heat-treated BLG in the presence of glucose, the response of macrophages was analysed by measuring the levels of secreted IL-8, CCL20 and IL-1β. As shown in Fig 8, non-processed native BLG and low-temperature dry-heated BLG induced significant levels of IL-8, CCL20 and IL-1β secretion in macrophages when compared to medium. High-temperature dry-heating of BLG significantly reduced the secretion of all three analysed cytokines when compared to native BLG. A similar reduction was observed in wet-heated BLG for IL-8 and IL-1β, but less strongly and not significantly for CCL20.

**Fig 8.**
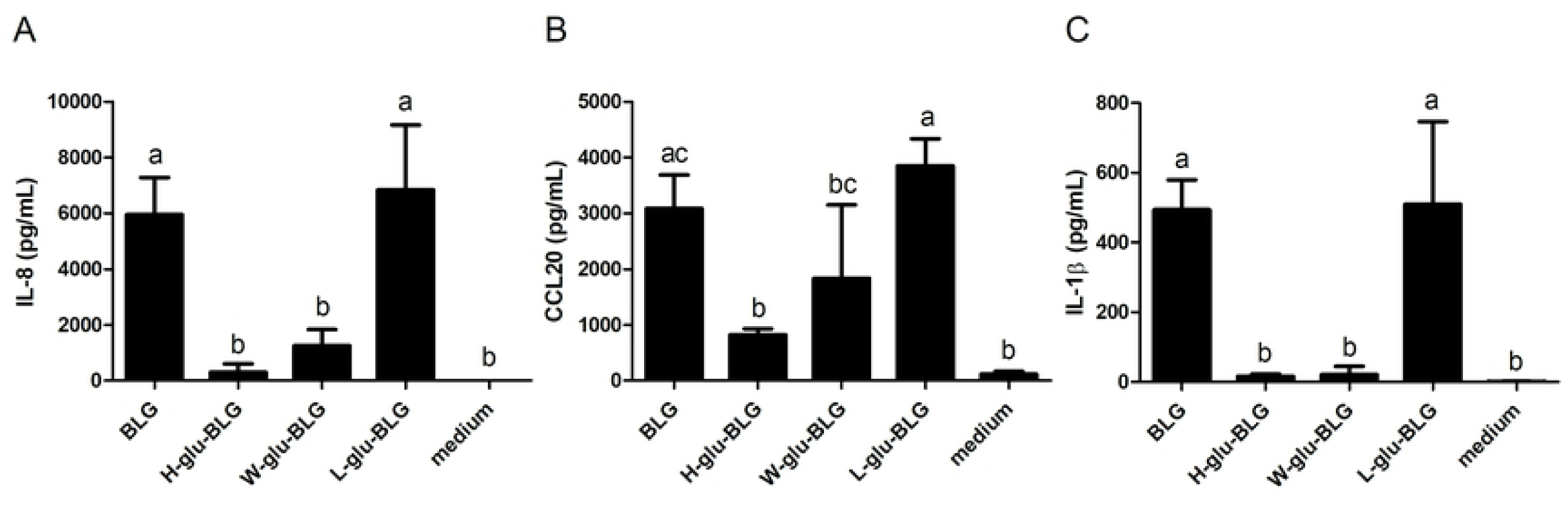
Cytokine and chemokine responses of macrophages upon exposure to native or heat-treated BLG varies. BLG was untreated or heated (W, H or L) in the presence of glucose and the soluble fraction was incubated with macrophages. The cytokine secretion of IL-8 (A), CCL20 (B) and IL-1β (C) was analysed in the supernatant with ELISA. The results represent mean values ± SD of 2-4 independent experimental measurements. The statistical differences were calculated with Tukey’s Test with different letters (a–c) in the bars indicated significantly different (p < 0.05).

## 4. Discussion

In an earlier study, we observed that hydrophobicity and the aggregation state of heat-processed BLG were important parameters that were related to the protein uptake by THP-1 macrophages [3]. Here, we extended these findings by analysing whether similar relations would apply for 1) a protein with similar pI but higher molecular weight (thyroglobulin); 2) a protein with similar molecular weight but higher pI (lysozyme); 3) EDC-aggregated proteins; and 4) another cell type by using THP-1 dendritic cells. The comparison between THP-1-derived macrophages and dendritic cells regarding uptake revealed similar patterns for the three tested proteins and the different heat-treatments. Macrophages and DCs are both able to capture exogenous particles and induce an immunogenic response, playing an essential role in immunological defence [18]. We did observe, however, a quantitative difference in uptake, as the uptake values for wet-heated thyroglobulin and BLG were relatively higher in macrophages than in DCs. This suggests that macrophages are more efficient in their phagocytic role than DCs which is in line with literature stating that uptake by macrophages is part of their core role in eliminating harmful substances, whereas uptake by DCs is a means to initiating an adaptive immune response [19]. Upon comparing the three tested proteins, we found that the physicochemical alterations upon heat-treatment to thyroglobulin and BLG were similar but different from lysozyme. These proteins differ in size (thyroglobulin (330 kDa) > BLG (18.4 kDa) ∼ lysozyme (14.3 kDa)), pI (lysozyme (10.7) > BLG (5.2) ∼ thyroglobulin (4.5)) and denaturation temperature (lysozyme (75 °C) > BLG (65 °C) ∼ thyroglobulin (60°C)) [20-23], suggesting that both pI and heat stability may be important for heat-induced structural protein changes. The high pI may have conferred a positive charge to lysozyme during wet-heating (pH 7.4), thereby hindering aggregation under this condition due to electrostatic repulsion. The strong internal coherence and stability of lysozyme’s structure may also have contributed to its stability during heat processing [24]. It is reported that the addition of the saccharides would impact the physicochemical characteristics, in particular in combination with the different applied heat-treatments [25]. But for the soluble fraction analysed, we found only a very limited influence of glycation on protein uptake.

Significant increases in uptake of thyroglobulin by macrophages correlated with increases in hydrophobicity and aggregation that occurred upon wet-heating (Fig 3 and 4), similar to BLG. This is in line with the findings on lysozyme, where the lack of increased uptake upon heating of lysozyme by macrophages may have been related to the lack of aggregation and only minimal increases in hydrophobicity upon heat-treatment. In order to better distinguish between aggregation and hydrophobicity as drivers of uptake, we used EDC to chemically crosslink proteins through the carboxyl and amino groups of amino acids (Fig 5), thereby being able to generate protein aggregates without applying a thermal treatment. Polymerisation was induced for BLG and thyroglobulin, as expected. Unexpectedly, EDC was unable to induce polymerisation in lysozyme, but it increased its hydrophobicity instead. This might be due to the intramolecular crosslinking of lysozyme between Lys-13 and Leu-129 as described [26], upon EDC treatment. In contrast to intermolecular crosslinking, which occurs for most proteins upon EDC treatment, the induction of intramolecular linkages might lead to protein unfolding resulting in exposure of hydrophobic regions. The strong increase in uptake of EDC-treated lysozyme indicates that hydrophobicity is a more important driver for uptake of heat-treated proteins than aggregation. A correlation between hydrophobicity of peptides and uptake by HeLa cells was also reported earlier [27]. Heating of proteins, through its resulting increased exposure of hydrophobicity, may thus underlie the increased uptake by APCs.

Next, we aimed to elucidate the uptake route of BLG samples by macrophages. Uptake in general can be categorized into clathrin-dependent and clathrin-independent. In clathrin-dependent uptake, also known as receptor-dependent uptake, cells recognise and internalize the receptor-bound substance [28] whereas clathrin-independent uptake functions via caveolae structures (membrane domains) or actin (taking up large particles up to 5 μm) [29]. Caveolae are reported to be involved in the uptake by APCs [30]. Caveolae-dependent uptake might be the basic route for the uptake of BLG by macrophages as inhibition of this route led to a reduced uptake for all samples (Fig 6). Uptake of BLG, that had been wet-heated in the presence of glucose, by macrophages seemed to involve other endocytosis routes (Fig 6C). The blocking of microphagocytosis led to a significant decrease in uptake of wet-heated and also of BLG that had been high-temperature dry-heated in the presence of glucose, probably due to the presence of higher molecular weight particles. The clathrin-dependent route might also be involved in the uptake by macrophages of BLG that had been wet-heated in the presence of glucose, as a significant decrease was found when the respective inhibitor was used. As shown in Fig 7, BLG wet-heated in the presence of glucose had clearly higher binding capacity for sRAGE, galectin-3 and CD36 compared to native BLG. Uptake mediated by the CD36 receptor was reported to be clathrin-dependent, while galectin-3 and RAGE are thought to be clathrin-independent [31-33]. However, it is doubtful if these receptors do play essential roles in the significantly enhanced uptake of BLG that had been wet-heated in the presence of glucose, as these receptors are mainly reported to be involved in the uptake of glycated proteins [34-36]. There was no strong glycation of BLG that had been wet-heated in the presence of glucose as shown in our previous study [3]. The binding of BLG, that had been wet-heated in the presence of glucose, to the receptors may be due to the high hydrophobicity of the sample. For example, the strong affinity of the S100 protein for RAGE was ascribed to a binding mechanism in which hydrophobic structures are involved [37, 38]. High-temperature dry-heated BLG, which had a high degree of glycation, bound to all receptors to a higher extent than wet-heated BLG, but its uptake was not significantly reduced when clathrin-dependent uptake was blocked. In another study, the uptake of BLG that had been heated in solution at 60 °C for 10 days with glucose by dendritic cells was reported to be RAGE independent but scavenger receptor dependent [39]. Scavenger receptor A was reported to be involved in the uptake of glycated ovalbumin by macrophages [40], which could thus be a potential target for future discrimination. The increased uptake of wet-heated BLG may thus be due to enhancing of the clathrin-/receptor-dependent route, although the involved receptors still need to be clarified.

IL-8, CCL20 and IL-1β are proinflammatory cytokines secreted by macrophages when they are activated by antigens [41]. However, there seems to be no direct relation between the macrophage’s cytokine release and its uptake of BLG. As shown in Fig 8, the cytokine production of macrophages treated with native and low-temperature dry-heated BLG was generally significantly higher than for the other samples. The strong cytokine release of macrophages in response to native BLG exposure might (partly) be related to the higher levels of LPS contamination in this sample, as shown in S1 Fig. Native BLG had an LPS concentration of 155 pg/mL, while wet-heated, low- and high-temperature dry-heated BLG in the presence of glucose contained 2.3, 7.8 or 4.6 pg/mL, respectively. A previous study demonstrated that 500 pg/mL LPS induced a 7-fold increase in IL-8 production by THP-1 macrophages when compared to medium [42]. Here, we found a production of 6,000 pg/mL of IL-8 by macrophages treated with BLG compared to no production when treated with medium. Thus, although the cytokine production of macrophages in response to BLG exposure might be partly endotoxin-driven, the BLG-induced IL-8, and also CCL20 and IL-1β, production is much higher than would be expected from the level of LPS contamination alone. When comparing the samples, the minor structural changes following low-temperature dry-heating of BLG did not change the levels of cytokine responses. In contrast, high-temperature dry-heated or wet-heated BLG, with significant structural changes compared to native BLG, only induced limited cytokine responses by macrophages. This is in line with findings from another study where BLG-mediated production of cytokines by murine bone-marrow derived dendritic cells was also reduced through heating of the BLG, which these authors claimed to be due to glycation. However, in this study BLG was heated in solution with glucose for 10 days at 60 °C, similar to our wet-heating process. This makes it likely that also in their samples, physicochemical changes such as increased hydrophobicity, rather than glycation, led to a lowered cytokine response [39]. Increased hydrophobicity of our BLG samples did not induce clear immunological responses but rather lowered these. Seong and Matzinger hypothesized hydrophobic structures (‘hyppos’) as molecular patterns that initiate immune responses (DAMPs), playing a role in aetiology of certain pathologies [43]. Although we found that hydrophobicity was strongly correlated to uptake of protein (aggregates), we could not establish a direct link with immunogenicity in our strictly *in vitro* experimental models. Considering the differences in types of proteins that were included in our study vs. the endogenously present, malformed proteins that are the basis of Seong and Matzinger’s hypothesis, it is perhaps too early to correlate our findings to the DAMP-concept.

## Conclusions

We found strong similarities in uptake of processed proteins with different molecular weight but similar pI (BLG vs THY), but different uptake kinetics for a processed protein with similar molecular weight but strongly differing pI (BLG vs LYS). Uptake of proteins by macrophages and dendritic cells showed strong qualitative similarities, indicating that the protein uptake process is, to a considerable extent, independent of the phenotype of APCs. The collective data on processed and EDC-treated thyroglobulin, lysozyme and BLG reveals that hydrophobicity is the most important determining factor for uptake. Various mechanisms appeared to be involved in the uptake of variously processed forms of BLG. Binding to receptors that are known to bind AGEs was strongest for more damaged forms of BLG, but production of number of cytokines by macrophages was strongest in response to native and minimally modified BLG. Nevertheless, the obtained data showed clearly that there are immunological consequences of physicochemical modifications due to different heating methods, which is of relevance to the food industry in developing new methods to minimize possible adverse immunological effects of processed food.

## Author contributions

YD acquired and analyzed all data. IL and IS contributed to the data collection. YD, CG, KH and HW participated in the design and interpretation of the reported experiments and results. YD drafted and edited the manuscript. CG, MT, KH and HW revised and edited the manuscript. All authors read and approved the final manuscript.

## Acknowledgments

This work was financial supported by the China Scholarship Council (No. 201507720020).

## Supporting information captions

**S1 Fig. There is no significant difference between the original uptake value of BLG, lysozyme and thyroglobulin for THP-1 derived macrophages (Mϕ, A) and immature dendritic cells (iDC, B)**. The result represents the mean value ± SD of 4 independent cell experiments. No significant differences have been found using unpaired T-test with Welch’s correction.

**S2 Fig. Wet-heating and high-temperature dry-heating significantly altered the physicochemical properties of native thyroglobulin**. Thyroglobulin (THY) was untreated or heated (W, H or L) in the absence or presence of saccharides (glu, lac or GOS) and a number of physiochemical parameters (i.e., solubility, loss of amino group, AGE-related fluorescence, hydrophobicity, α-helix and β-sheet structure) were measured. The results represent mean values ± SD of n=4 measurements of 2 independent experiments based on 2 independent sample sets. Statistical differences compared to native THY were calculated with Dunnett’s Test: *p < 0.05; **p < 0.01; ***p < 0.001. ND: not detectable.

**S3 Fig. Aggregation of thyroglobulin was prominent upon wet-heating and high-temperature dry-heating in the presence of GOS**. Thyroglobulin (THY) was untreated or heated (W, H or L) in the absence or presence of saccharides (glu, lac or GOS) and aggregation in the soluble fraction was measured using size exclusion chromatography. The data points represent the average values of 2 independent sample sets.

**S4 Fig. Significant physicochemical modifications of native lysozyme were mainly induced by high-temperature dry-heating**. Lysozyme (LYS) was untreated or heated (W, H or L) in the absence or presence of saccharides (glu, lac or GOS) and a number of physicochemical parameters (i.e., solubility, loss of amino group, AGE-related fluorescence, hydrophobicity, α-helix and β-sheet structure) were measured. The results represent the mean values ± SD of n=4 measurements of 2 independent experiments based on 2 independent sample sets. Statistical differences compared to native LYS were calculated with Dunnett’s Test: *p < 0.05; **p < 0.01; ***p < 0.001.

**S5 Fig. Wet-heated lysozyme samples had physicochemical properties that are similar to native lysozyme**. Lysozyme (LYS) was untreated or heated (W, H or L) in the absence or presence of saccharides (glu, lac or GOS) and tested for solubility, uptake by THP-1 macrophages (uptake), AGE formation (AGE), glycation (OPA), percentage of α-helix (helix) or β-sheet (sheet), and exposure of hydrophobic regions (ANS). The aggregation-related parameters, proportion of monomer, oligomers and polymers are not shown as they did not differ between the samples.

